# Regulatory T cells control type 1-driven immunopathology restraining GM-CSF-producing helper T cells

**DOI:** 10.1101/2024.01.24.577048

**Authors:** Sara Costa-Pereira, Margit Lanzinger, Myrto Andreadou, Nicolas Nunez, Juan Villar-Vesga, Francesco Prisco, Philipp Häne, Elsa Roussel, Sinduya Krishnarajah, Rachel Chanel Lindemann, Frederike Westermann, Laura Oberbichler, Aakriti Sethi, André Fonseca Da Silva, Mirjam Lutz, Sonia Tugues, Sarah Mundt, Anja Kipar, Melanie Greter, Donatella De Feo, Burkhard Becher

## Abstract

Regulatory T (T_reg_) cells are critical for maintaining peripheral tolerance and preventing autoimmunity. T_reg_ cell depletion or dysfunction rapidly results in fatal multiorgan inflammation linked to unrestrained effector T cell expansion, but the cytokine network underlying immunopathology, and its direct cellular mediators, remain elusive. Here, we combined gene targeting, fate-mapping tools, and high-dimensional cytometry to identify the T helper (T_H_) cell-derived cytokines and responding cells that execute inflammatory tissue damage upon global loss of peripheral tolerance in mice. We found that T_H_ cell-derived GM-CSF, but not IL-17A, directed the ensuing immunopathology and thereby mortality through recruitment of tissue-invading phagocytes and granulocytes, and enhancement of their reactive oxygen species production and phagocytic proficiency. Our study highlights the critical role of T_reg_ cells in controlling GM-CSF- producing T_H_ cells and type 1-responses to restrain phagocyte-mediated tissue destruction and provides a framework for the use of anti-GM-CSF therapies in patients with chronic inflammatory disorders.

## INTROUCTION

Autoimmune diseases will affect approximately 10% of the world population, and their prevalence is increasing^1^. These conditions arise when parts of the immune system attack and destroy otherwise normal and healthy tissues^2^. Despite thymic negative selection, peripheral autoreactive T cells are found in healthy individuals and considered the main drivers of autoimmunity^3–5^. This highlights a strong need for peripheral tolerance, a delicate and fine-tuned process in which regulatory T (T_reg_) cells are the central players, at once controlling self-reactive T cell clones^6^, restraining inflammation, and limiting immunopathology^7^. The fundamental role of T_reg_ cells is well illustrated by the effects of mutations in the gene encoding *Foxp3*, the transcription factor that induces and sustains T_reg_ cell functionality^8,9^. In humans, *Foxp3* mutations cause severe immune dysregulation, polyendocrinopathy, enteropathy, X-linked (IPEX) syndrome^10^, in which multisystemic autoimmune inflammation causes death in the first years of life^11^. Immunosuppression and allogeneic hematopoietic stem cell transplantation are partially effective strategies, but prognosis remains bleak for these patients and insights into better therapeutic options are much needed^12^.

Scurfy mice, which bear a frameshift mutation in *Foxp3*, likewise suffer extreme immunopathology and so serve as a model for IPEX^13^. Similarly, adult mice depleted of *Foxp3*^+^ T_reg_ cells develop uncontrolled lympho- and myelo-proliferative disease accompanied by a cytokine storm that ultimately leads to fatal multi-organ failure^14^. Dysregulated CD4^+^ T_H_ cells are the key cell type driving inflammation in this setting, superseding the contribution of CD8^+^ T cells^15^. Through diverse mechanisms, T_reg_ cells largely, directly and indirectly, restrain self-reactive CD4^+^ T_H_ cells in peripheral tissues to protect against aberrant immune responses (review in ^16^). In the context of the global loss of tolerance triggered by T_reg_ cell depletion, cytokine storms are one of the most prominent hallmarks, with T_H_ cells secreting high amounts of pro-inflammatory factors^14,17^.

In the realm of T_H_ cell-mediated autoimmune chronic inflammation, both type 1 and type 3 polarization states^18^ have been implicated, with GM-CSF and IL-17, respectively, commonly recognized as the primary cytokines accountable for orchestrating the inflammatory response. GM-CSF and IL17A accumulation at sites of tissue inflammation has been reported in atopic dermatitis and psoriasis lesions^19–22^, the synovial fluid of patients with rheumatoid arthritis (RA)^23,24^ and the cerebrospinal fluid of patients with multiple sclerosis (MS) during the active^25^ and the relapsing-remitting^26^ phase of disease. Both cytokines are also capable of inducing severe graft-versus-host-disease (GVHD) in allogenic recipients^27,28^. Of note, in experimental autoimmune encephalomyelitis (EAE) -a well-established model for autoimmune tissue inflammation - auto-aggressive GM-CSF-producing T_H_ cells have a non-redundant role in immunophatology^29–31^, whilst the contribution of IL-17A to EAE is a matter of debate^32–36^. Notably, a role of T_reg_ cells in controlling GM-CSF- and IL-17A- producing T_H_ cells has been suggested in psoriasis^37^ and intestinal inflammation^38^. A major open question pertains to the nature of the T_H_ cell response and its cytokine profile kept in check during homeostasis by T_reg_ cells to prevent autoimmunity, and which responder cells orchestrate widespread tissue damage upon global loss of peripheral tolerance.

Here, we present a systematic immune profiling of secondary lymphoid and immune-damaged tissues in mice depleted of T_reg_ cells. We reveal the T_H_-derived instructive cytokine network and the responding cells executing tissue damage, and uncover new facets to the critical role of GM-CSF during immunopathology triggered by the global loss of peripheral tolerance.

## RESULTS

### Loss of T_reg_ cells leads to GM-CSF^+^ T_eff_ cell expansion and tissue infiltration by inflammatory phagocytes

To investigate the effects of the loss of T_reg_ cell-mediated immune suppression, we conducted a longitudinal profiling of *Foxp3*^DTR^ mice^14^ in which injection of diphtheria toxin (DT) causes global ablation of *Foxp3*^+^ T_reg_ cells (Figure 1A). Here, DT-treated *Foxp3*^DTR^ mice suffered diffuse dermatitis with epidermal ulcers and crusts, broncho interstitial pneumonia with severe bronchus-associated lymphoid tissue expansion, and almost complete loss of pancreatic Langerhans islets: all lesions were surrounded by extensive inflammatory infiltrate, and mice began to die from 10 days post-DT (dpDT) injection (Figure S1A). To analyse the lymphoid and myeloid cell populations present we designed a high-dimensional spectral flow cytometry [HD-Cyto] antibody panel and applied it to the spleen and axillary lymph nodes (Ax LNs) and the three most-affected tissues (lung, skin and pancreas) (Figure 1A). This revealed a complete depletion of *Foxp3*^+^ T cells from tissues by 5 dpDT (Figure 1B). Using uniform manifold approximation and projection (UMAP)-dimensionality reduction and FlowSOM-guided clustering, we identified the lymphoid cell clusters present (Figures. S1B and S1C). By day 10 dpDT, mice had approximately double the number of effector CD4^+^ and CD8^+^ T cells in lymphoid and target tissues compared to day 0, and all tissues except for the skin also showed expansion of B cells; this coincided with reduced numbers of naïve T cells in the spleen and Ax LNs of T_reg_-cell-depleted mice (Figure 1C). Among innate immune cells, we found myeloid, NK, and γδ T cells increased in all analysed organs at 10 dpDT, while innate lymphoid cell (ILC) numbers were unaffected (Figure 1C). Trends towards these changes were also evident by 5 dpDT (Figure S1D).

**Figure 1.**
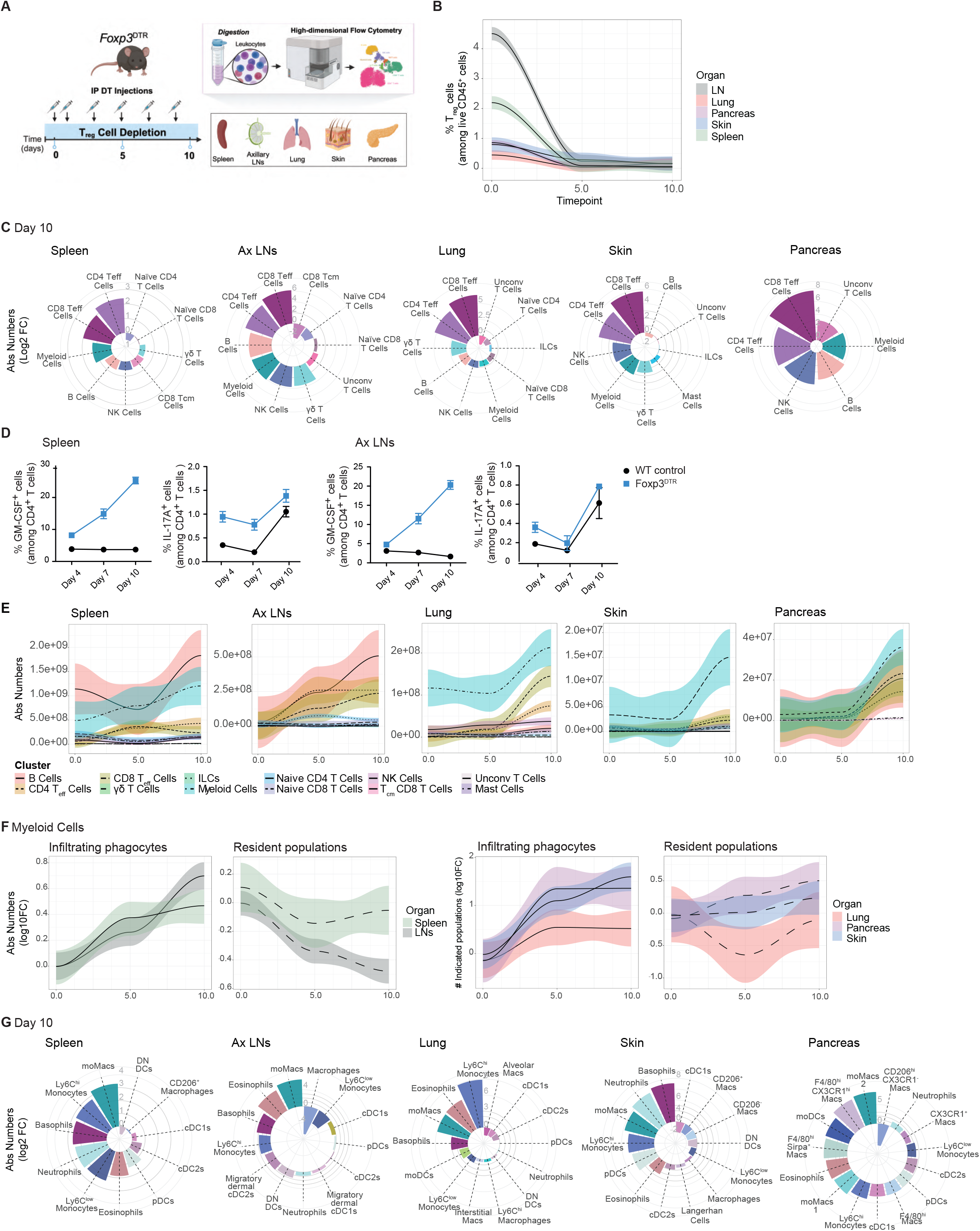
Loss of T_reg_ cells leads to the expansion of T_eff_ cells and myeloid cell infiltration in lymphoid and target organs. (A) Experimental workflow (B) Line plot depicting the frequency of T_reg_ cells at 0, 5 and 10 dpDT in indicated tissues (n=3 mice per timepoint, data from one experiment, mean shown as black centerline ± s.d. (coloured area)). (C) Absolute numbers of the identified cell populations in the spleen, Ax LNs, lung, skin and pancreas of *Foxp3*^DTR^ mice at 10 dpDT. Cell numbers are indicated as log2 fold change from day 0 (n=3 mice per timepoint, data from one experiment). (D) Cytokine production among CD4^+^ T Cells in the spleen and Ax LNs of DT-treated WT control (black) or Foxp3^DTR^ (blue) mice (n=3 mice per timepoint, data from one experiment). (E) Absolute numbers of the identified leukocyte populations at 0, 5 and 10 dpDT in the indicated lymphoid and target tissues (n=3 mice per timepoint, data from one experiment, mean shown as black dashed centerline ± s.(D) (coloured area)). (F) Absolute numbers of infiltrating phagocytes and resident myeloid cell populations at 0, 5 and 10 dpDT in indicated tissues. Cell numbers shown as log10 fold change from day 0 (n=3 mice per timepoint, data from one experiment, mean shown as black centerline ± s.d. (coloured area)). (G) Absolute numbers of the identified cell populations in the spleen, Ax LNs, lung, skin and pancreas of *Foxp3*^DTR^ mice at 10 dpDT. Cell numbers are indicated as log2 fold change from day 0 (n=3 mice per timepoint, data from one experiment).

We next measured the impact of T_reg_ cell depletion on the production of the type 1 and type 3 cytokines GM-CSF and IL-17A, respectively. We uncovered increasing frequencies of GM-CSF^+^ CD4^+^ T_H_ cells in the spleen and Ax LNs during DT treatment, while IL-17A^+^ T_H_ cells remained rare throughout (Figure 1D).

Type 1 cytokines, such as GM-CSF, primarily influence myeloid cells, which are implicated in several chronic inflammatory and autoimmune disorders^39,40^. Here, myeloid cells were the dominant infiltrate across target organs, surpassing even the expanded T_eff_ cell population (Figure 1E); infiltrating phagocytes predominated, while the resident compartment was reduced or unchanged in all organs apart from the pancreas (Figure 1F). FlowSOM-guided analysis of the CD45^+^ CD90^-^ myeloid fraction across all organs identified: neutrophils (Ly6G^+^), eosinophils (Siglec-F^+^) and basophils (CD11b^+^CD88^+^); Ly6C^hi^ (Ly6C^high^CD64^+^) and Ly6C^low^ (Ly6C^int/low^CX3CR1^high^PD-L1^high^) monocytes and monocyte-derived macrophages (moMacs) (Ly6C^high^MHCII^high^); as well as conventional dendritic cells (cDCs) types 1 (XCR1^high^CD11b^-^) and 2 (CD11b^high^CD172a^high^), and plasmacytoid dendritic cells (pDCs) (CD11b^-^ MHCII^+^CD38^+^Ly6C^high^) (Figures S1F and S1G). We also identified tissue-resident and inflammatory macrophage subsets based on their expression of CD11b, F4/80, CD64, CD206, CX3CR1, SIRPα, MHCII, CD38 and Siglec-F (Figures S1E and S1F). Compared to day 0, at 10 dpDT, we saw an increase in Ly6C^hi^ monocytes, moMacs, neutrophils and eosinophils; this was most pronounced for Ly6C^hi^ monocytes in the lung (465-fold increase). Baseline myeloid populations, including cDCs and tissue-resident macrophages, contracted upon T_reg_ cell depletion in most organs, aside from the spleen and pancreas, while pDCs were increased in the spleen, skin and pancreas (Figure 1G), potentially recruited there by monocyte-derived cells^41^. Alongside, both inflammatory infiltrating and resident macrophages were expanded in the pancreas (Figure 1G). That these changes were in-progress at 5 dpDT (Figure S1G), suggests rapid mobilization of the myeloid compartment into the target tissues upon T_reg_ cell depletion. Immunofluorescent imaging of lung and pancreata from DT-treated mice confirmed the predominance of Iba1^+^ mononuclear phagocytes (MNPs) and revealed their proximity to T cells (Figure S1H).

In conclusion, the loss of T_reg_-mediated tolerance in these mice induced unbridled T cell expansion and tissue invasion, a shift from naïve to effector T cell phenotypes, and increasing predominance of GM-CSF production over time, concomitant with marked infiltration of inflammatory phagocytes.

### GM-CSF is the main driver of tissue inflammation upon loss of peripheral tolerance

The tissue invasion of phagocytes upon T_reg_ cell depletion suggested a dysregulated type 1 immune response, with little evidence for the involvement of type 3 mechanisms. To confirm this hypothesis, we crossed *Foxp3*^DTR^ mice with mice deficient in either the canonical type 1 cytokine GM-CSF (*Csf2*^-/-^) or its type 3 counterpart, IL-17A (*Il17a*^-/-^). While IL-17A deficiency did not ameliorate fatal immunopathology caused by T_reg_ cell depletion, GM-CSF-deficiency conferred a 50% increase in survival and significantly lower histology scores in the skin, lung and pancreas, compared to GM-CSF-sufficient *Foxp3*^DTR^ mice (Figure 2A-D). The splenomegaly and increased ear thickness characteristic of *Foxp3*^DTR^ mice were also partially rescued by the loss of GM-CSF (Fig 2E).

**Figure 2.**
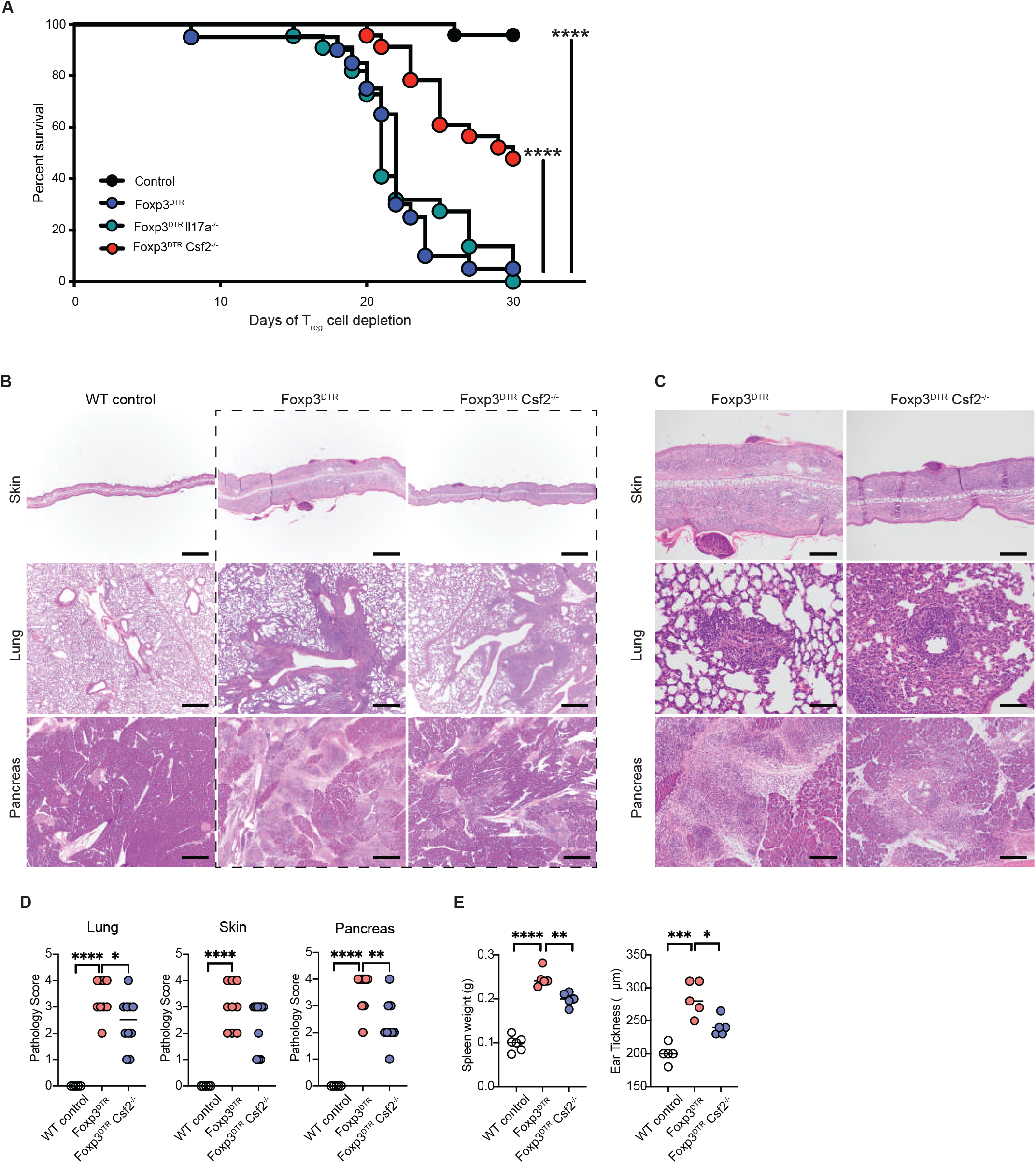
GM-CSF is the main driver of tissue inflammation upon loss of peripheral tolerance. **(A)** Kaplan-Meier-survival curve of DT-treated WT control (black, n=24), *Foxp3*^DTR^ (blue, n=20), *Foxp3*^DTR^*Csf2*^-/-^ (red, n=18), and *Foxp3*^DTR^*Il17a*^-/-^ (turquoise, n=22) mice (Data from two independent experiments, Mantel–Cox test; ****P < 0.0001). **(B)** Representative H&E-stained sections of indicated tissues from DT-treated WT control, *Foxp3*^DTR^ and *Foxp3*^DTR^*Csf2*^-/-^ mice at 18 dpDT (Images representative of two independent experiments). **(C)** Representative H&E-stained sections of indicated tissues *Foxp3*^DTR^ and *Foxp3*^DTR^*Csf2*^-/-^ mice of panels in (B) with higher magnification (40X). **(D)** Pathological inflammation scores of indicated tissues from WT control (white), *Foxp3*^DTR^ (red) and *Foxp3*^DTR^*Csf2*^-/-^ (blue) mice at 18 dpDT (n=5-9 mice per timepoint, data from two independent experiments, t-test; *P < 0.05; **P < 0.005; ****P < 0.0001). **(E)** Spleen weight and ear thickness in WT control (white), *Foxp3*^DTR^ (red) and *Foxp3*^DTR^*Csf2*^-/-^ (blue) mice at 14 dpDT (n=5 mice per timepoint, data from one independent experiment, t-test; *P < 0.05; **P < 0.005; ***P < 0.0005; ****P < 0.0001).

Altogether, we identified a dominant role of GM-CSF in orchestrating widespread immunopathology induced by loss of peripheral tolerance.

### Fate-mapping identifies T_mem_ cells as the main source of GM-CSF upon Treg cell depletion

To unequivocally identify the cellular sources of GM-CSF *in vivo* upon T_reg_ cell depletion, we crossed *Foxp3*^DTR^ mice with FROG^Ai14^ fate-mapping mice^31^, in which GM-CSF*-*expressing cells (tdTomato^+^ EGFP^+^), as well as their progeny and other cells that have previously produced GM-CSF (tdTomato^+^) are fluorescently labelled. We then characterised the labelled cells in lymphoid and target organs by HD-Cyto (Figure 3A). This revealed increased numbers of tdTomato^+^ and tdTomato^+^ EGFP^+^ cells across all tissues during T_reg_ cell depletion (Figure 3B and S2A), predominantly from among the CD45^+^ fraction (Figure S2B). In the lung, 2.6% of GM-CSF production stemmed from non-immune cells, which we assume to be epithelial cells^42^ (Figure S2B).

**Figure 3.**
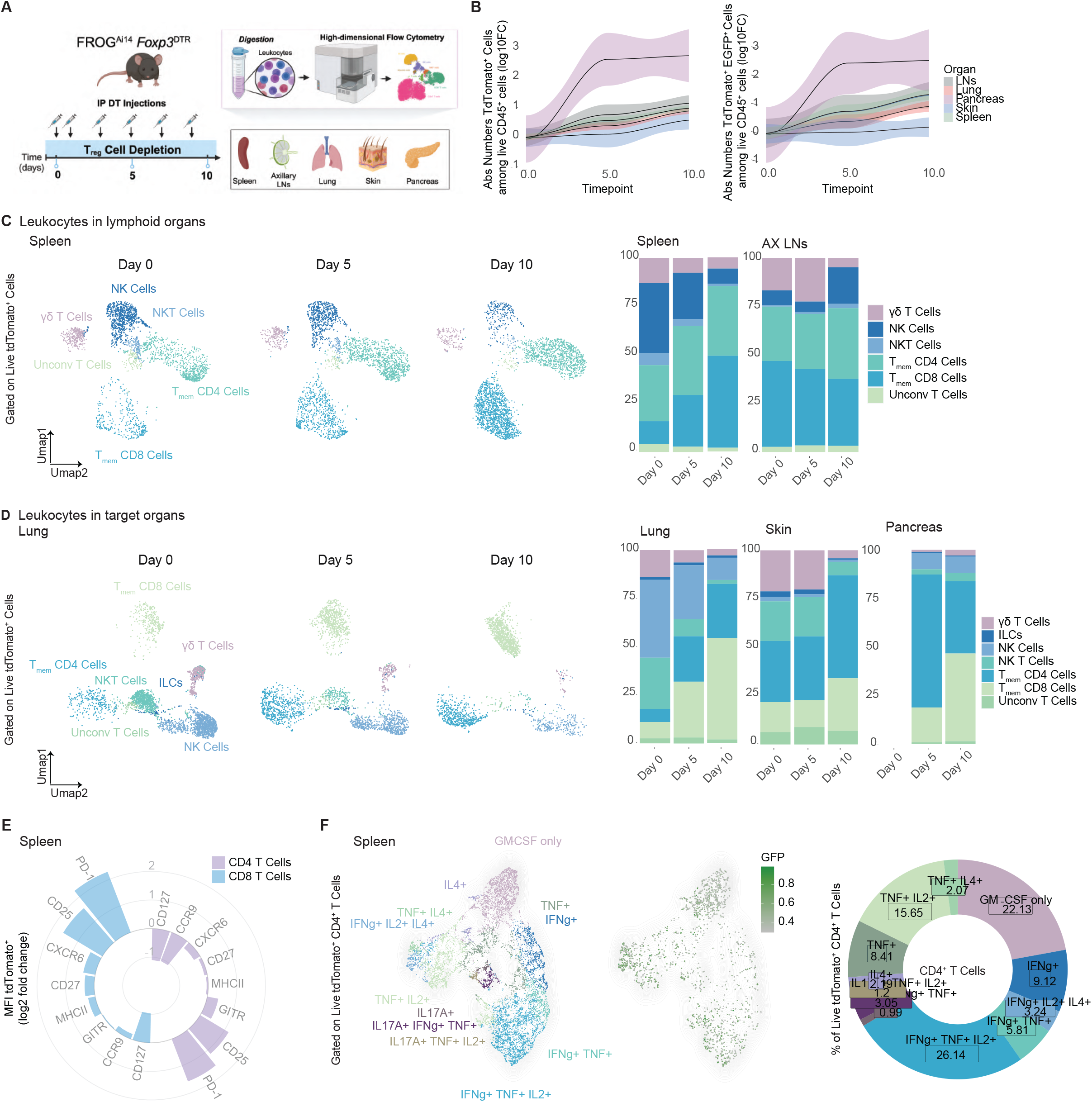
Fate-mapping identifies T_mem_ cells as the main source of GM-CSF upon Treg cell depletion. **(A)** Experimental workflow. **(B)** Number of live CD45^+^ tdTomato^+^ and live CD45^+^ tdTomato^+^ GFP^+^ cells at 0, 5 and 10 dpDT in indicated tissues from FROG^Ai14^*Foxp3*^DTR^ mice. Cell numbers are indicated as log10 fold change from day 0 (n=2-3 mice per timepoint, data are representative of two independent experiments, mean shown as black centerline ± s.(D) (coloured area)). **(C)** UMAP displaying 20,000 randomly sampled CD45^+^ tdTomato^+^ cells from the spleens of DT-treated mice at the indicated timepoints. On the right, bar graphs display the population frequencies among CD45^+^ tdTomato^+^ cells in each lymphoid organ at the indicated timepoints (n=2-3 mice per timepoint, data are representative of two independent experiments). **(D)** UMAP displaying 20,000 randomly sampled CD45^+^ tdTomato^+^ cells from the lungs of DT-treated mice at the indicated timepoints. On the right, bar graphs display the population frequencies among CD45^+^ tdTomato^+^ cells in each target organ at the indicated timepoints (n=2-3 mice per timepoint, data are representative of two independent experiments). **(E)** Expression profile of CD4^+^ and CD8^+^ tdTomato^+^ T cells from the spleen of FROG^Ai14^*Foxp3*^DTR^ mice at 10 dpDT, shown as log2 fold change from day 0 (n=2 mice per timepoint, data are from one experiment). **(F)** UMAP displaying 5000 randomly sampled tdTomato^+^ CD4^+^ T cells isolated from the spleen of FROG^Ai14^*Foxp3*^DTR^ mice at 10 dpDT. On the right a UMAP shows the EGFP^+^ cells overlay, and a doughnut chart displays the quantification of the identified cytokine-producing populations among tdTomato^+^ CD4^+^ T Cells (n=2 mice per timepoint, data from one experiment).

In lymphoid organs at baseline, the tdTomato^+^ cell population comprised T memory (T_mem_) CD4^+^ and CD8^+^ cells, unconventional T cells, and γδ T cells, with NK and NK T cells contributing to the splenic GM-CSF^+^ pool; however, by 10 dpDT CD4^+^ T_mem_ and CD8^+^ T_mem_ cells had become the dominant source of GM-CSF in both organs (Figure 3C). In target tissues, by contrast, 50-70% of labelled cells at baseline were innate: in the lung, unconventional T cells and NKT cells were the predominant sources of GM-CSF, while in the skin, 50% of GM-CSF-producing cells were γδ T cells, ILCs, NK and NKT cells (Figure 3D). Of note, there was no detectable tdTomato signal in the pancreas at baseline (Figure 3D). Following T_reg_ cell depletion, most organs exhibited increased dominance of GM-CSF-producing T cells, with this effect being most pronounced in the spleen and skin (Figure 3D).

When we compared tdTomato^+^ cells at days 0 and 10 dpDT, we saw increased expression of programmed cell death protein 1 (PD-1) and CD25, with reduced CD127 expression, suggesting a terminally differentiated effector state (Figure 3E). We also detected increased CXCR6 expression, particularly in labelled cells from the lung (Figure S2C), consistent with a resident cytotoxic T_H_ cell phenotype that is associated with prolific cytokine production^43,44^. Direct assessment of the cytokine-production pattern of these cells in the spleen revealed that GM-CSF expression either occurred alone, or accompanied by expression of IFN-γ and/or TNF and IL-2; a minor population of cells also co-expressed IL-17A (Figure 3F).

Altogether, while at baseline innate lymphoid cells contributed significantly to GM-CSF production, upon loss of peripheral tolerance fully differentiated T_H_ cells became the major source of GM-CSF, which was mostly co-expressed with the other type 1 cytokines IFN-γ and TNF.

### GM-CSF-sensing phagocytes execute tissue damage through oxidative burst and phagocytosis

GM-CSF is an important communication conduit between pathogenic T cells and tissue-invading phagocytes during chronic inflammatory conditions and autoimmunity (reviewed in ^45,46^). To dissect the contribution of GM-CSF to tissue immunopathology in the absence of T_reg_ cells, we compared the changes in the immune compartment of *Foxp3*^DTR^*Csf2*^-/-^ mice and *Foxp3*^DTR^ mice upon T_reg_ cell depletion. To minimize the confounding factors steaming from different states of tissue damage between the two genotypes that could influence cell isolation, we analysed cells at 5 dpDT - the earliest timepoint of effective T_reg_ cell depletion (Figure 1B). The size of the T, B and NK cell compartments was not widely affected by GM-CSF and T_reg_ cell deficiency at this timepoint (Figure S3A). However, phagocyte infiltration into target organs during T_reg_ cell depletion was relatively less in *Foxp3*^DTR^*Csf2*^-/-^ compared to *Foxp3*^DTR^ mice (Figure 4A). Our data, in line with previous reports^47,48^, identifies GM-CSF as a recruiter of neutrophils and inflammatory phagocytes into sites of tissue destruction. Accordingly, we also found more granulocytes and monocytes in the spleen of *Foxp3*^DTR^*Csf2*^-/-^ mice (Figure S3B), suggesting impaired egress and recruitment into tissues.

**Figure 4.**
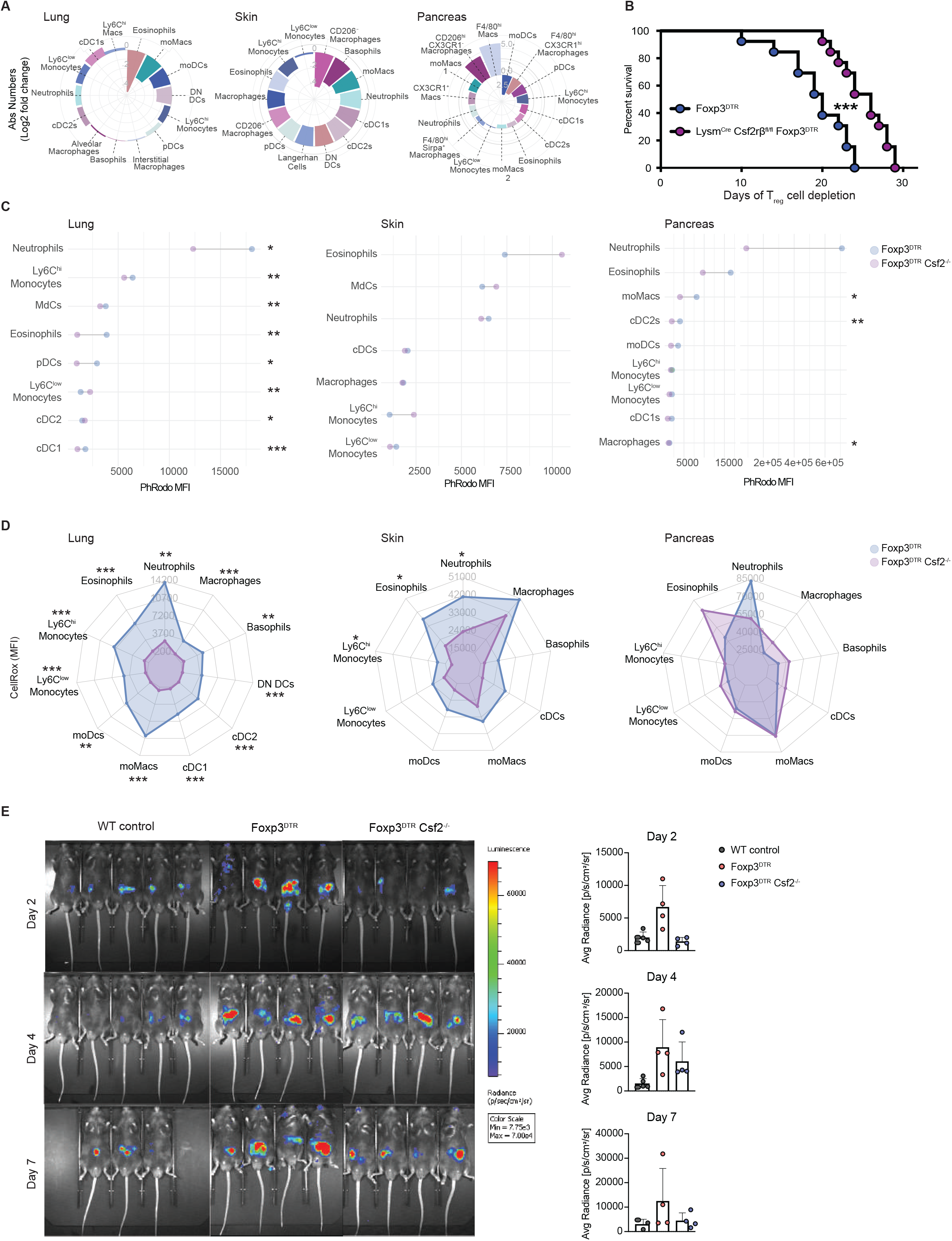
GM-CSF-sensing phagocytes execute tissue damage through oxidative burst and phagocytosis. **(A)** Absolute numbers of the identified populations in the lung, skin and pancreas of *Foxp3*^DTR^*Csf2*^-/-^ mice, shown as normalized population cell numbers at 5 dpDT compared to day 0, between *Foxp3*^DTR^*Csf2*^-/-^ and *Foxp3*^DTR^ mice, then plotted as the relative log2 fold change (n=3 mice per timepoint, data from one experiment). **(B)** Kaplan-Meier-survival curve of DT-treated *Foxp3*^DTR^ mice (blue, n=13) and *LysM*^Cre^C*sf2rb*^fl/fl^*Foxp3*^DTR^ (purple, n=13) mice (data from two independent experiments, Mantel– Cox test; ***P < 0.001). **(C)** Median fluorescence intensity (MFI) of internalized red pHrodo E. coli bioparticles in the indicated populations from the lung, skin and pancreas of *Foxp3*^DTR^ and *Foxp3*^DTR^*Csf2*^-/-^ mice at 8 dpDT (n=5 mice per timepoint, data are representative of two independent experiments, t-test; *P < 0.05; **P < 0.01; ***P < 0.001). **(D)** MFI of the fluorogenic probe CellRox among the indicated populations isolated from the lung, skin and pancreas of *Foxp3*^DTR^ and *Foxp3*^DTR^*Csf2*^-/-^ mice at 8 dpDT (n=3-5 mice per timepoint, data are representative of two independent experiments, t-test; *P < 0.05; **P < 0.01; ***P < 0.001). **(E)** *In vivo* imaging of ROS/RNS post-L-012 chemiluminescent probe injection into WT control, *Foxp3*^DTR^ and *Foxp3*^DTR^*Csf2*^-/-^ mice on 2, 4 and 7 dpDT. On the right side, bar plots depict the quantification of the L-012 average radiance signal measured as photons/s/cm^2^/sr. (n=4-5 mice per timepoint, data are representative of two independent experiments).

To confirm that GM-CSF-sensing cells were the main executors of tissue immunopathology, we crossed *Foxp3*^DTR^ with *LysM*^Cre^*Csf2rb*^fl/fl^; this allowed us to deplete T_reg_ cells while ablating GM-CSF signalling in granulocyte/monocyte/macrophage/DC lineages^49^. Accordingly, among GM-CSF-stimulated splenocytes from these mice, phosphorylation of the signal transducer and activator of transcription (STAT)5 induced by GM-CSF receptor engagement was significantly reduced in neutrophils, monocytes and macrophages, and partially reduced in cDCs (Figure S3C). Notably, T_reg_ cell depletion in *LysM*^Cre^*Csf2rb*^fl/fl^*Foxp3*^DTR^ resulted in a significant increase in overall survival compared to sex- and age-matched *Foxp3*^DTR^ controls (Figure 4B).

We next identified the main tissue-damaging effector mechanisms of polymorphonuclear phagocytes (PMPs) and MNPs in the presence and absence of GM-CSF. Monocyte-derived cells and neutrophils in the skin of *Foxp3*^DTR^*Csf2*^-/-^ mice displayed a lower median expression of TNF compared to *Foxp3*^DTR^ (Figure S3D), while monocyte-derived cells in the lung and pancreas exhibited lower phagocytic capacity (Figure 4C). Alongside, we observed GM-CSF-dependent reactive oxygen species (ROS) production in phagocytes from lung and skin at 8 dpDT (Figure 4D). *In vivo* imaging confirmed the higher accumulation of ROS and reactive nitrogen species (RNS) upon T_reg_ cell depletion in *Foxp3*^DTR^ mice compared to DT-treated WT controls, which was evident as early as 2 dpDT. Consistently, *Foxp3*^DTR^*Csf2*^-/-^ mice showed reduced ROS/RNS accumulation early upon T_reg_ cell loss (Figure 4E).

Altogether these findings support the dominant role of GM-CSF signalling in controlling the pathogenic program of PMPs and MNPs during tissue inflammation and immunopathology following the global loss of peripheral tolerance mediated by T_reg_ cells.

## DISCUSSION

T_reg_ cells maintain peripheral tolerance, restrain inflammation and limit immunopathology^7^. Previous studies identified a T_H_ cell-derived cytokine storm as the main executor of tissue damage in scurfy mice^50^ and IPEX patients^51^, with both type 1 and type 3 as likely culprits of initiating tissue inflammation^52^. Our goal here was to dissect the T_H_ cell-derived cytokine network controlled by T_reg_ cells and to understand the mechanisms involved in tissue destruction and organ failure during loss of peripheral tolerance. Leveraging *Foxp3*^DTR^ mice as a valuable tool for investigating T_reg_ cell steady-state functions in adult mice, we found that T_H_ cell-derived GM-CSF, but not IL-17A, was necessary for immunopathology. In the absence of GM-CSF, phagocyte invasion was dramatically reduced, highlighting its role in communication between pathogenic T cells and tissue-invading phagocytes^45^. Consistently, upon T_reg_ cell depletion, tissue-invading myeloid cells outnumbered lymphocytes in affected tissues: detailed analysis identified Ly6C^hi^ monocytes, monocyte-derived phagocytes, and granulocytes among the increased populations, while numbers of resident macrophages and cDC declined. In line with these findings, Hu et al. observed that, upon reconstitution of the depleted T_reg_ cell pool, the control of myeloproliferation was dependent on the suppression of activated T cells^7^. We confirmed the relevance of GM-CSF- sensing across tissue-invading phagocytes by showing improved survival of *LysM*^Cre^*Csf2rb*^fl/fl^*Foxp3*^DTR^ mice upon T_reg_ cell depletion. Nevertheless, we cannot formally exclude other GM-CSF-sensing cell types – most notably, non-immune cells – as contributors to GM-CSF-mediated pathology. T cells, on the other hand, do not express the GM-CSFR and are not directly affected by the presence or absence of GM-CSF^27,45,53^.

The critical effector properties of myeloid cells that confer protection against pathogens makes them equitably likely to drive tissue damage. GM-CSF is a powerful modulator of inflammatory phagocytes’ effector functions in diverse disease settings^27,47,54^. Mechanistically, in the absence of peripheral tolerance, T_H_-cell-derived GM-CSF drives increased ROS production and phagocytosis across tissue-infiltrating inflammatory phagocytes and granulocytes. Interestingly, we found organ-specific differences in these phenomena, suggesting that GM-CSF support of phagocyte’ effector functions is strongly influenced by the surrounding microenvironment. For instance, neutrophils and moMacs that accumulated in the skin upon T_reg_ depletion showed GM-CSF-dependent increases in ROS production but not phagocytosis; while in the pancreas, GM-CSF strongly enhanced the phagocytic ability of inflammatory moMacs. These findings parallel our previous work, where genetically-induced GM-CSF dysregulation led to an influx of monocytes and granulocytes, with organ-specific immunopathological consequences^55^. Therefore, the relative contributions of inflammatory phagocytes and their effector functions to immunopathology appears to strongly depend on the local tissue microenvironment.

We also demonstrated that T_H_-cell derived GM-CSF production and the type 1-response are the main targets of peripheral tolerance. The central role of GM-CSF in orchestrating immunopathology has been well illustrated in pre-clinical models of chronic inflammatory diseases, such as MS^47^, RA^56^, psoriasis^19^ and GVHD^27^. Several studies have reported impaired immunosuppressive functions in the T_reg_ cell compartment in these disorders, and adoptive transfer of functional T_reg_ cells is considered an attractive immunotherapeutic approach^57^, despite being technically challenging. Preclinical and clinical investigations suggest that T_reg_ cells could prevent GVHD without impeding the graft-versus-tumour effect in the context of hematopoietic cell transplantation (HCT)^58–60^; likewise, anti-GM-CSF treatment limits GvHD without compromising alloreactive T cell control of the tumour^27^ and its safety and efficacy its currently being assessed in a randomized clinical trial (ISRCTN42585840).

In conclusion, we show that control of pathogenic GM-CSF-producing T_H_ cells is the predominant mechanism used by peripheral T_reg_ cells to prevent tissue inflammation. Specifically, GM-CSF orchestration of granulocytes’ and inflammatory phagocytes’ chemotaxis and effector functions dictates the tissue destruction observed upon loss of peripheral tolerance. This demonstrates a specific suppression of type 1-immune responses by T_reg_ cells to prevent catastrophic immune activation. Further studies will be important to dissect the self-tolerance mechanisms employed by T_reg_ cells to curb this pathogenic T_H_ cell subset. Our findings offer a foundational basis and support for the emerging efforts to employ anti-GM-CSF therapies in a wide range of autoimmune and chronic inflammatory conditions.

### Limitations of the study

Whilst we have uncovered a dominant role of GM-CSF in immunopathology induced by the absence of T_reg_ cells, our data do not preclude the effects of other tissue-damaging pathways controlled by T_reg_ cells. For example, B cells contribute to inflammatory tissue infiltration and decreased survival in scurfy mice^61^; this could result from the emergence of IL-4, as seen after depletion of T_reg_ cells in *Foxp3*^DTR^ mice^17^, to which B cells are strong responders^62^. However, how B cell expansion fits into the inflammatory cascade warrants further investigation. We also found that GM-CSF-producing T_H_ cells co-expressed other cytokines, most prominently IFNγ and TNF. The role of IFNγ in chronic inflammation, for instance, is highly complex. While it has strong pro-inflammatory features and effects on various cell types, including NK cells and phagocytes, powerful immunosuppressive properties have also been described across various models of chronic inflammation^63–66^. The positions of these mediators in the inflammatory cascade initiated by T_reg_ cell depletion should be addressed in future studies.

## MATERIALS AND METHODS

### Mice

Animal experiments were approved by the Swiss Veterinary Office and were performed according to federal and institutional guidelines. All mice were kept in individually ventilated cages under specific-pathogen-free conditions on a 12h light/dark cycle with free access to water and chow diet. C57BL/6 (WT) mice were obtained from Janvier. *Foxp3*^DTR-GFP^ mice (referred to as *Foxp3*^DTR^) were generated by Kim et al.^14^, kindly provided by Markus Feuerer (Deutsches Krebsforschungszentrum, Heidelberg, Germany) and backcrossed to the CD90.1 background. Mice deficient in GM-CSF (*csf2^-/-^*) were kindly provided by Glenn Dranoff^67^ and backcrossed into the C57BL/6 background for 9 generations. Yoichiro Iwakura (University of Tokyo, Japan) provided *il17a^-/-^* mice^68^. *LysM*^Cre^ mice were obtained from Björn Clausen (Universitätsmedizin Mainz, Germany)^49^ and were kept heterozygous*. Csf2rb^fl^*mice were generated by crossing *Csf2rb^LacZ^* mice^47^ to a flp-deleter strain bred in house. Experiments were conducted with either male or female mice aged 6-14 weeks. Control and experimental animals were age- and sex-matched in each experiment.

### *In vivo* treatments

T_reg_ cell-depletion was achieved by intraperitoneal (IP) injection 500 ng diphtheria toxin (DT) (Merck-Millipore) dissolved in of 150 µl of PBS on days 0 and 1, and subsequently every other day during the indicated time frame. For survival experiments, DT injections were performed on days 0 and 1 and subsequently three times per week.

### Generation of single-cell suspensions

Following euthanasia by CO_2_ inhalation, mice were perfused with approximately 20 ml of ice-cold PBS before organ removal. For myeloid cell studies, Ax LNs, spleen, lungs, pancreas and skin (ear) were collected. The samples were cut into small pieces and digested in HBSS containing 10% (v/v) fetal calf serum (FCS), 100 µg/ml deoxyribonuclease (DNase) I (Sigma-Aldrich) and one of the following enzyme solutions: 0.2 mg/ml Collagenase type IV (Sigma-Aldrich) for 30 min (LNs and spleen); 0.4 mg/ml Collagenase type IV for 45 min (Lung) at 37°C; 1 mg/ml Collagenase type IV for 15 min (Pancreas) at 37°C. Ear skin samples were first divided into dorsal and ventral sides, then digested in 0.25 mg/ml Liberase (Roche, 5401119002) and 200U/ml DNase I (Sigma, DN25-100MG) for 20 min at 37°C before mechanical disruption using a gentleMACS Octo dissociator. After digestion, all tissues were disrupted by passage through an 18-gauge needle, then filtered through a 70 µm- or 100µm- pore diameter cell strainer and washed with PBS. Red blood cells were removed from the lung and spleen cell suspensions by incubation with 1 ml of red blood cell lysis buffer (4.15 g NH4Cl, 0.55 g KHCO3 and 0.185 g EDTA in 500 ml ddH_2_O) for 2 min on ice and subsequent washing with PBS. For exclusive T cell studies, T cells from Ax LNs and spleen were isolated by mechanical disruption using a 70μm cell strainer and washed with PBS.

### Flow cytometry

Flow cytometric analysis was performed following standard methods. For surface labelling, cells were incubated with the respective antibodies for 25 min at 4°C and washed with PBS, followed by a 20 min streptavidin staining step. When applicable, chemokine staining was achieved by a 20 min incubation at 37°C. Cell fixation and permeabilization was performed using the Foxp3/Transcription Factor Staining Buffer (eBioscience) kit according to the manufacturer’s instructions. For intracellular cytokine detection, cells were restimulated ex vivo for 4-5h at 37 °C in complete RPMI medium (RPMI supplemented with 2mM L-Glutamine and 10% fetal calf serum (FCS)) with GolgiPlug (BD, 1:1,000 dilution), phorbol 12-myristate 13-acetate (50ng/ml, Applichem) and Ionomycin (500ng/ml, Invitrogen). Cells were fixed and permeabilized using the BD Cytofix/Cytoperm™ solution (BD) kit according to the manufacturer’s instructions. Cells were incubated with the respective antibodies overnight at 4°C for intracellular labelling and washed with perm buffer (BD). Fluorescence data were acquired on a Cytek Aurora using SpectroFlo v2 Software, and analysed using FlowJo v10 (TreeStar). For pSTAT5 staining detection, cells were firstly surface stained with antibodies conjugated to methanol stable fluorochromes. Subsequently, cells were stimulated for 45 min with complete medium supplemented or nor with 20 ng/ml GM-CSF at 37°C. Cells were then fixed with 2 % PFA for 20 min at 4°C, washed and permeabilized with ice-cold methanol for 20 min at 4°C. Lastly, cell were stained with an anti-pSTAT5 antibody and antibodies conjugated to methanol sensitive fluorochromes. For all experiments, dead cells were excluded from the analysis using Near-IR Live/Dead fixable staining reagent (BioLegend). Doublets were excluded by FSC-Area versus FSC-Height gating.

### Histology

Following euthanasia by CO_2_ asphyxiation, mice underwent transcardial perfusion with ice-cold PBS before tissue harvesting and fixation by incubation in 10% neutral buffered formalin between 12-48h at 4°C. Tissue dehydration was performed in 70% ethanol. Samples were embedded in paraffin and 3µm-thick tissue sections were cut on a microtome (Microm HM 360, Thermo Fisher) and transfered to glass slides. Paraffin-embedded sections were subject to haematoxylin and eosin (H&E) staining and analysed using an Eclipse Ci-L microscope (Nikon) using 10x or 40x objectives. The histological assessment was undertaken in a blinded fashion.

Inflammatory changes were quantified based on the adaption of the scoring system previously published^69^, as follows: Score 0, no inflammatory changes were evident in the examined sections; Score 1, minimal inflammation characterized by the sparse presence of lymphocytes and plasma cells around bronchioles in the lung, and multifocally in the pancreas and skin samples; Score 2, mild inflammation characterized by slightly increased numbers of lymphocytes and plasma cells and occasional neutrophils around bronchioles in the lung, and multifocally in the pancreas and skin samples; Score 3, moderate inflammation characterized by increased quantities of lymphocytes, plasma cells, neutrophils and scattered areas of more severe inflammation or necrosis. These areas were observed around bronchi and bronchioles in the lung and multifocally in the pancreatic interstitium and skin dermis, where they were associated with multifocal shallow erosions or a few small ulcerations in the skin; Score 4, marked inflammation characterized by multifocal to coalescing or diffuse infiltrates composed of abundant degenerate and nondegenerate neutrophils, lymphocytes and plasma cells associated with variable areas of necrosis and cellular debris. These areas surrounded bronchi and bronchioles in the lung and were multifocal in the pancreatic interstitium and skin dermis, associated with multifocal erosions to locally extensive ulcerations covered by a serocellular crust.

### Immunofluorescence microscopy

Following death by CO_2_ asphyxiation, mice underwent transcardial perfusion with approximately 20ml of ice-cold PBS followed by 10ml of 4% paraformaldehyde in PBS (pH 7.2). After immediate tissue removal, fixation was carried out by overnight incubation in 4% paraformaldehyde with PBS at 4°C. Upon PBS washing, the tissue was transferred to 30% sucrose in PBS for 24–72 h at 4°C, placed into cryo-embedding medium (Medite, 413011-00) and frozen at −20°C. 14-μm-thick sections were cut using a Cryostat (CryoStar NX70) onto glass slides. Samples were incubated in working solution (1% Triton X-100, 1% FCS and 1% Donkey Serum (SUPP)) with primary antibodies 12-48h at 4°C. The following day, samples were washed in PBS containing 1µl/ml Tween 20% followed by a 30min incubation with secondary Alexa Fluor-conjugated antibodies and DAPI in the working solution. Sections were washed with PBS and, when required, incubated 6h with the secondary antibody staining in the working solution. After mounting in anti-fading mounting medium and sealed under a coverslip images were acquired on a Zeiss LSM 980 Airyscan inverted confocal laser scanning microscope or a Leica Stellaris 5 upright using 10x or 40x oil immersion objective and analysed with Fiji software (NIH, Bethesda).

### Phagocytosis assay

Single-cell suspensions from the lung, skin and pancreas of DT-treated mice were suspended in 200µl of complete RPMI medium containing 4µl of pHrodo *E. coli* BioParticles (Thermo Fisher) after surface staining. Cells were incubated for 90 minutes at 37 °C, transferred to ice and fluorescence data immediately acquired on a Cytek Aurora (SpectroFlo v3.0).

### *In vivo* imaging of reactive oxygen and nitrogen species

Mice were subjected to IP injection of 100 µl of the luminescent probe L-012 (Wako Chemical) dissolved in ultrapure H_2_O (25 mg/kg). Mice were anesthetized with isoflurane and the bioluminescent signal was measured after 10 min using the IVIS spectrum *in vivo* imaging system with 2 min exposure time.

### High-dimensional data analysis

Data obtained from flow cytometry analyses underwent compensation, doublet and dead cell exclusion, pre-gating on CD45-positive and CD90-negative cells on FlowJo (TreeStar) before export. Using the R environment, a 0-1 normalization of each marker expression was performed. Unbiased analysis conducted on these data enabled dimension reduction through UMAP projections of stochastically selected cells^70^. Manual annotation of clusters based on the MFI of different markers was achieved using the FlowSOM algorithm (FlowSOM R package^71^). All plots were drawn using ggplot2. The primary R script was executed according to the instructions outlined in reference ^72^.

### Statistical analysis

The figure legends provide specific details on the experimental design and statistical tests conducted. Each mouse experimental group comprised 2-5 mice, with no exclusions of animals or data points. When applicable, data points were polled from all replicate experiments as specified in the figure legends. Statistical analyses were performed using Prism 9 (GraphPad Software) or Rstudio (version 4.1.0). P values were abbreviated as follows: p < 0.05 = *, p < 0.01 = **, p < 0.001 = ***, p < 0.0001 = ****, with only significant values being reported. Blinding was performed when possible, and all readouts are objective and not subject to experimental bias.

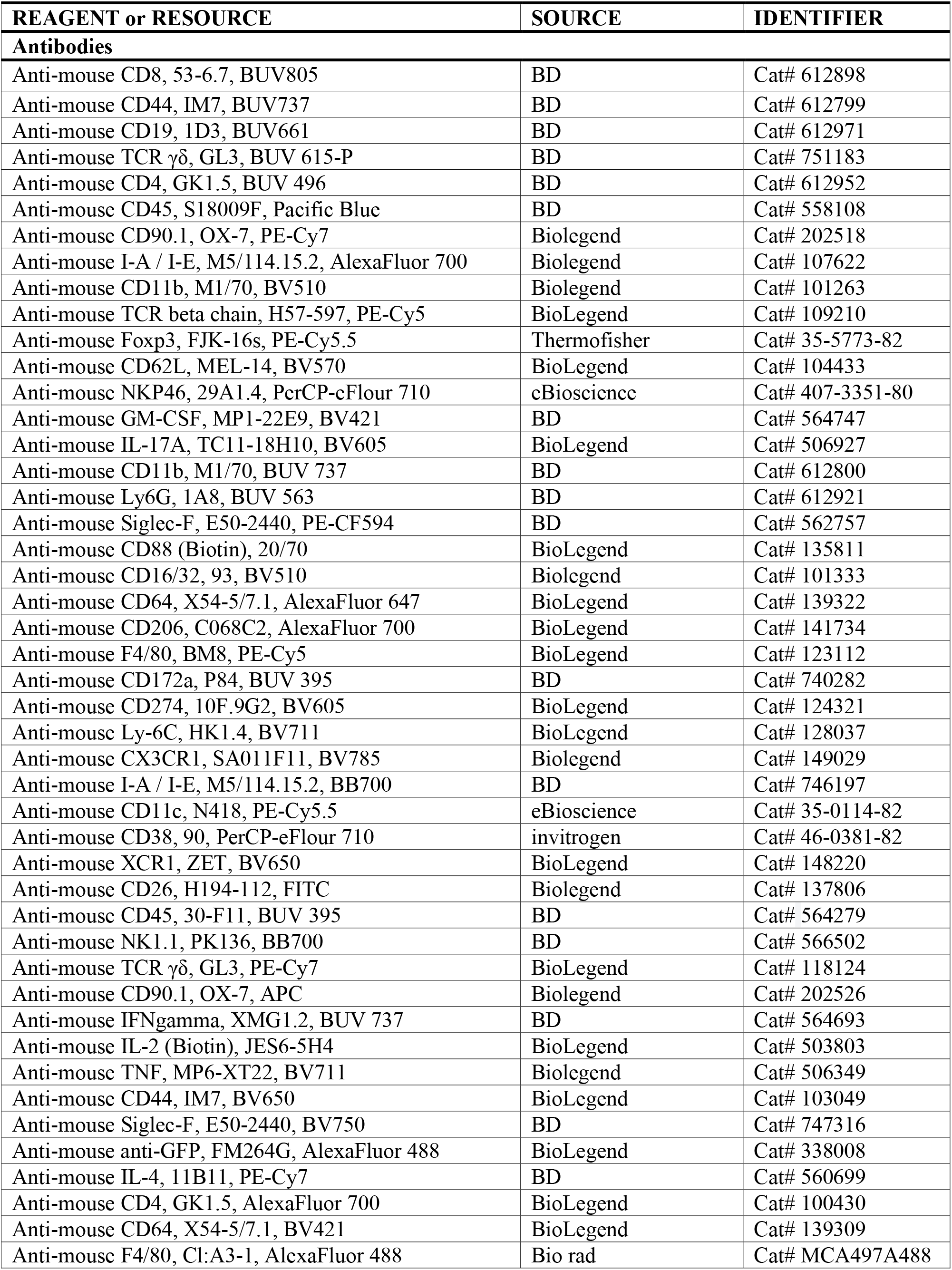
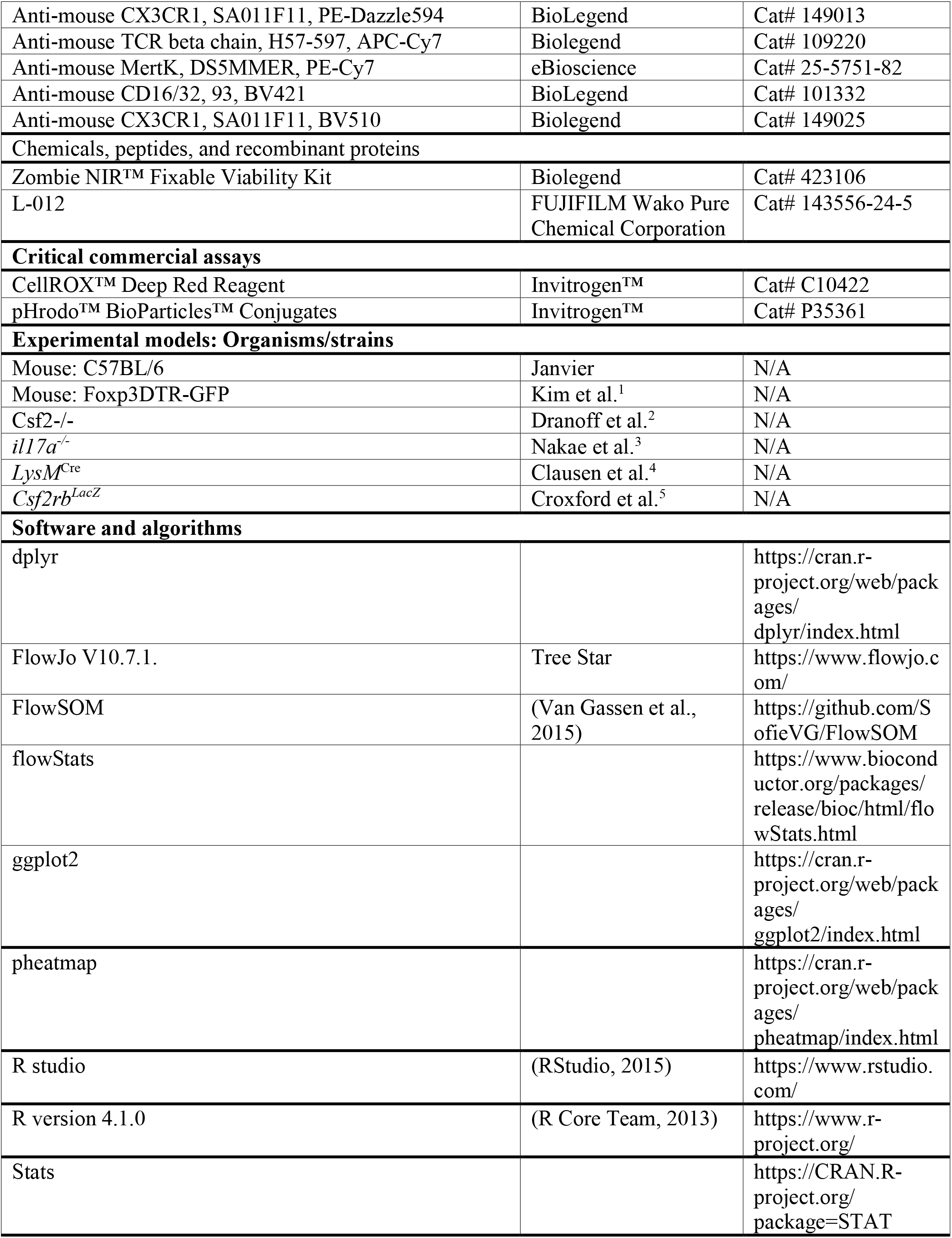
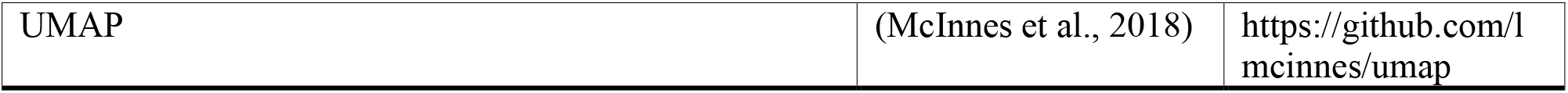

### Data and code availability

Data and code will be available upon request.

## ACKNOWLEDGMENTS

This work was supported by the Swiss National Science Foundation (310030_188450, and 310030_219287) to BB and the European Research Council (ERC; 882424) to B.B. D.D.F. received funding through: Forschungskredit Postdoc fellowship University of Zurich grant number: K-41302-10-01; Swiss Multiple Sclerosis Society (D.D.F., project: Regulation of phagocyte-mediated tissue damage in neuroinflammation and to D.D.F. project: F-41302-43 01. Progressive MS: disentangling the contribution of aging and chronic inflammation mechanisms to phagocyte-mediated neurodegeneration); through Kurt und Senta Herrmann Foundation. We express our gratitude to Dr. Lucy Robinson of Insight Editing London for the meticulous review and editing of the manuscript.

## AUTHOR CONTRIBUTIONS

Conceptualization, S. Costa-Pereira, M.L., M.A., D.D.F and B.B.; Methodology, S. Costa-Pereira, M.L., M.A., D.D.F, N.N., V.V.J., F.P., P.H., R.C.L., S.K., F.W., E.R., L.O., A.S., A.F.S. and M.L.; Formal Analysis, S. Costa-Pereira, M.L., F.P., N.N., and D.D.F.; Investigation, S. Costa-Pereira, M.L., F.P., M.A., N.N., and D.D.F.; Resources, B.B., A.K., S.M., M.G.; Writing – Original Draft, S. Costa-Pereira, D.D.F. and B.B.; Writing – Review & Editing, M.L., M.A., N.N., V.V.J., F.P., P.H., R.C.L., S.K., F.W., E.R., L.O., A.S., A.F.S., M.L., A.K., S.M., M.G., D.D.F., and B.B.; Visualization, S. Costa-Pereira, N.N., and D.D.F.; Funding Acquisition and Supervision, D.D.F. and B.B.

---

**Figure S1.**
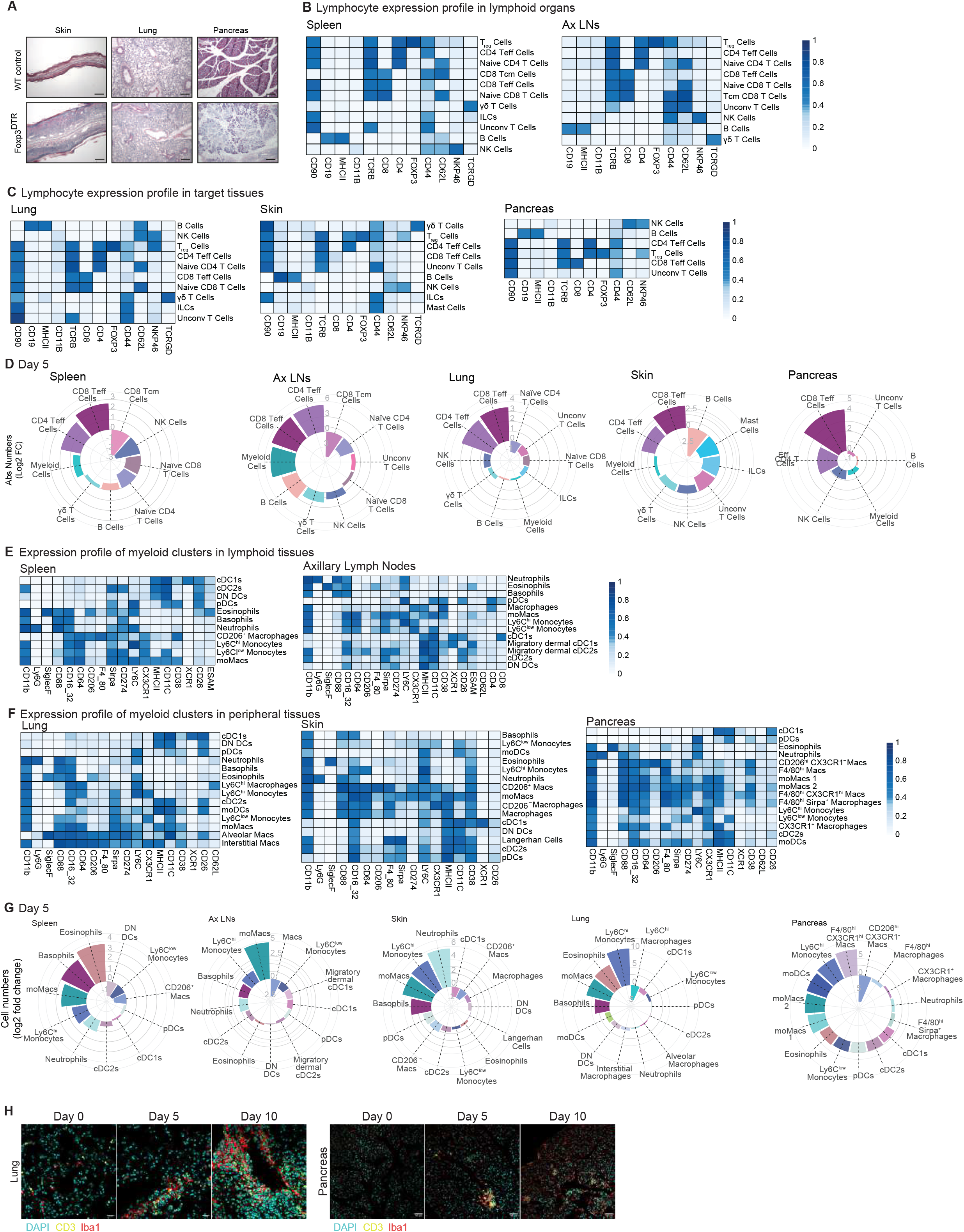
T_reg_ cell-depletion results in early infiltration of lymphoid and myeloid cells into lymphoid and target tissues. **(A)** Representative H&E staining of indicated tissues from DT-treated WT control and *Foxp3*^DTR^ mice at 18 dpDT. **(B)** Heatmap depicting the median marker expression of the identified lymphoid subsets from the lymphoid organs of T_reg_ cell-depleted mice. **(C)** Heatmap depicting the median marker expression of the identified lymphoid subsets from the target tissues of *Foxp3*^DTR^ T_reg_ cell-depleted mice. **(D)** Absolute numbers of the identified populations in the spleen, Ax LNs, lung, skin and pancreas of *Foxp3*^DTR^ mice at 5 dpDT. Cell numbers are indicated as a log2 fold change from day 0 (n=3 mice per timepoint, data are from one experiment). **(E)** Heatmap depicting the median marker expression of the identified myeloid subsets from the lymphoid organs of T_reg_ cell-depleted mice. **(F)** Heatmap depicting the median marker expression of the identified myeloid subsets from the target tissues of *Foxp3*^DTR^ T_reg_ cell-depleted mice. **(G)** Absolute numbers of the identified populations in the spleen, Ax LNs, lung, skin and pancreas of *Foxp3*^DTR^ mice at 5 dpDT. Cell numbers are indicated as log2 fold change from day 0 (n=3 mice per timepoint, data from one independent experiment). **(H)** Immunofluorescence microscopy showing DAPI (cyan), anti-CD3 (yellow) and anti-Iba1 (red) labelling of T_reg_ cell-depleted mice pancreas and lung at the indicated timepoints.

**Figure S2.**
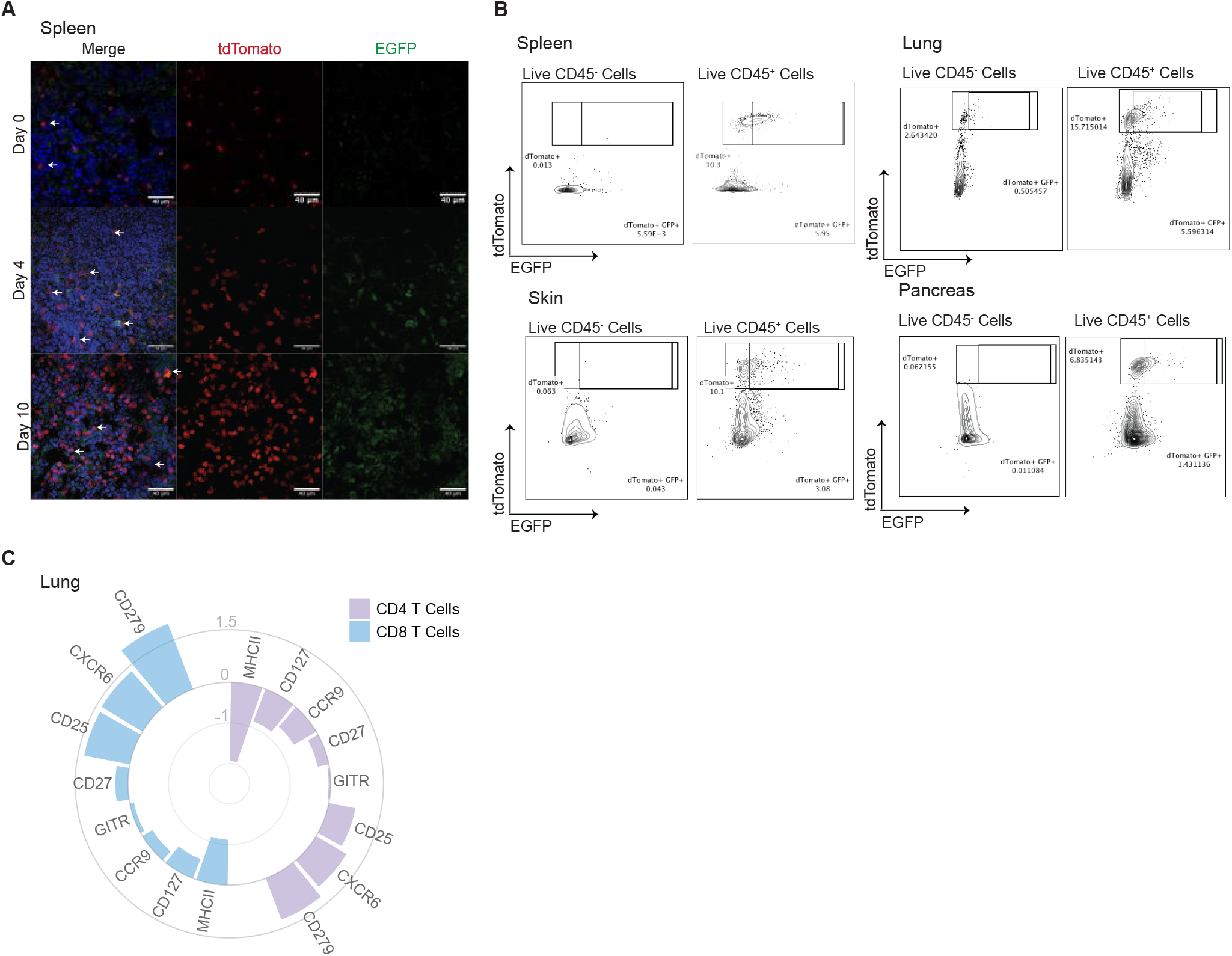
CD45^+^ compartment is the source of GM-CSF upon T_reg_ cell-depletion. **(A)** Immunofluorescence microscopy showing DAPI (Blue), tdTomato (Red) and EGFP (Green) in FROG^Ai14^*Foxp3*^DTR^ mice at 0, 5 and 10 dpDT. **(B)** Representative plots depicting the frequency of tdTomato^+^ EGFP^+^ cells among live CD45^-^ and CD45^+^ cells in the indicated organs. **(C)** Expression profile of CD4^+^ and CD8^+^ tdTomato^+^ T cells in the lung of FROG^Ai14^*Foxp3*^DTR^ mice at 10 dpDT shown as log2 fold change from day 0 dpDT (n=2 mice per timepoint, data from one independent experiment).

**Figure S3.**
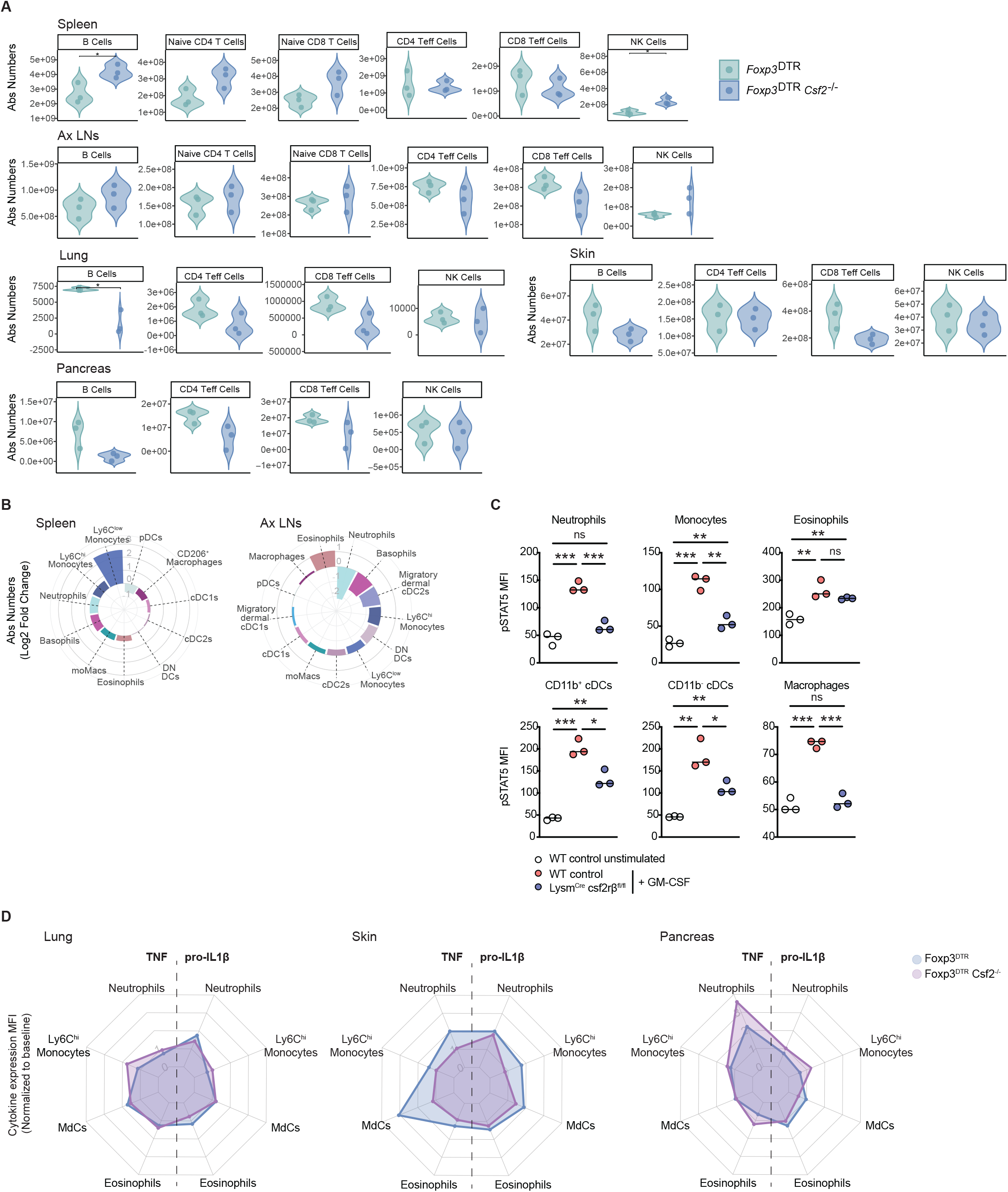
PMPs and MNPs are the GM-CSF-responsive cell types inducing chronic tissue inflammation upon the global loss of tolerance. **(A)** Absolute numbers of the identified populations in the indicated organs at day 5 upon T_reg_ cell- depletion, isolated from DT-treated *Foxp3*^DTR^ and *Foxp3*^DTR^*Csf2*^-/-^ mice. **(B)** Absolute numbers of the identified populations in the spleen and Ax LNs of *Foxp3*^DTR^*Csf2*^-/-^ mice indicated as normalized population cell numbers at 5 dpDT over day 0, between *Foxp3*^DTR^*Csf2*^-/-^ and *Foxp3*^DTR^ and then plotted as the relative log2 fold change (n=3 mice per timepoint, data from one independent experiment). **(C)** Individual pSTAT5 MFI values of the indicated populations from unstimulated WT control (white), stimulated WT control (red) or *LysM*^Cre^C*sf2rb*^fl/fl^ (blue) splenocytes (n=3 mice per timepoint, data are representative of two independent experiments, t-test, *P < 0.05; **P < 0.01; ***P < 0.001). **(D)** Cytokine expression among the identified myeloid cell subsets isolated from the lung, skin and pancreas of *Foxp3*^DTR^ and *Foxp3*^DTR^*Csf2*^-/-^ mice at 10 dpDT (n=3 mice per timepoint, data from one experiment).

